# Musical complexity governs a tradeoff between reliability and dimensionality in the neural code

**DOI:** 10.1101/2025.08.04.668380

**Authors:** Pauline G. Mouawad, Shievanie Sabesan, Alinka E. Greasley, Nicholas A. Lesica

## Abstract

The rich experience of listening to music depends on the neural integration of its constituent elements within the early auditory pathway. Here, we performed the first large-scale study of neural responses to complex music to characterize the neural coding of individual instruments and mixtures with both normal hearing and mild-to-moderate hearing loss. Using coherence and manifold analyses along with deep learning, we identified strong nonlinear interactions in mixed music that impacted the fidelity and geometry of the neural code. We found that increasing musical complexity resulted in the creation of new neural modes, but this increased dimensionality was associated with decreased reliability. This tradeoff persisted even after hearing loss, the effects of which were largely corrected with suitable amplification. These results suggest that the neural coding of music is governed by an inherent tradeoff and highlight a fundamental challenge in maintaining fidelity while processing sensory inputs with increasing complexity.

## Introduction

Music has always been fundamental to the human experience, but with the advent of modern digital technologies it is now ubiquitous. Many people spend much of their time listening to music [24], typically for enjoyment but also for social, cognitive, or emotional reasons [41, 25]. Music has emerged as a pivotal tool in cognitive science research; it is unique because it links sensory processing with reward, emotion, and arousal akin to physiological stimuli like food or sex [3, 48], yet more tractable for controlled lab investigation. Extensive research has investigated the power of music in communicating and eliciting affective responses [21, 30] and has enabled the development of computational models to characterize emotional expression in music [33]. Music also provides a natural framework for studies of statistical learning involving the construction and violation of sensory expectations [14], as well as experience-dependent plasticity [17].

Our ability to understand the higher-level processing of music [26, 34, 45, 16] is constrained by our limited knowledge of the neural code on which it relies: the music representation formed in the early auditory pathway. For example, the computation required to identify emotionally-salient musical features depends on the specifics of how those features are encoded in the periphery and the interplay between cortical and subcortical stages [29]. In contrast to the neural coding of speech, which has been extensively characterized from cochlea to cortex [47], the neural coding of complex music has never been studied using high-resolution neural recordings. Neural responses to simplistic music-like sounds or short excerpts have been analyzed to assess the processing of specific features (e.g., rhythm) [37, 38], but a comprehensive analysis of the neural responses elicited by complex music has not yet been undertaken.

Popular music from genres such as rock or pop is typically composed of many instruments – vocals, lead guitar, bass guitar, drums, keyboard, etc. – each of which has distinct acoustic properties. Because auditory processing is highly nonlinear, the neural representation of such music will reflect complex interactions between the individual instruments that impact the listening experience. Studies of musical scene analysis have demonstrated that the perception of individual instruments within mixed music depends on the specific instruments involved as well as their relative intensities [22, 42]. Furthermore, music perception involves both sensory and cognitive processes that are difficult to disentangle [23], especially for older listeners or those with hearing loss for whom the interplay between degraded input and altered brain dynamics can be particularly complex [40, 19, 4].

To characterize the sensory basis for music perception and how it is impacted by hearing loss, we made large-scale intracranial recordings from gerbils, which are commonly used as a model of human hearing because of their excellent sensitivity to low sound frequencies. We focused on the inferior colliculus (IC), the midbrain hub of the central auditory pathway where inputs from multiple brainstem pathways converge to form the neural representation that provides the basis for subsequent higher-level processing. We presented music of varying complexity and characterized the neural code at both the unit and network level. We also compared the neural code in animals with normal hearing to that in animals with mild-to-moderate hearing loss resulting from noise exposure. Our results reveal strong relationships linking musical complexity to the fidelity and geometry of the neural code that persist even after hearing loss.

## Results

We recorded neural activity from the IC of anesthetized gerbils using electrode arrays with a total of 512 channels spanning both hemispheres [1, 39]. We recorded activity in response to music from the *musdb18* database [36], which contains music in stem form with isolated stems for bass, drums, vocals, and other (all remaining instruments, e.g., guitar, keyboard). We used approximately 100 minutes of pop and rock songs, and presented each of the isolated stems as well as the full mixture in which the four stems were summed together. We processed the recordings to extract multi-unit activity (MUA) spike counts for each recording site, using 1.3 ms time bins to account for the fact that neurons in the IC can encode acoustic information with millisecond temporal precision [13].

### The basic properties of neural responses vary widely across music types

Example neural activity recorded from an animal with normal hearing in response to each of the 5 music types (the 4 stems – bass, drums, vocals, and other – and the mixture) is shown in Fig. 1a. The different music types vary widely in their spectrotemporal properties and this variation is reflected in the corresponding neural activity. The bass (B) and drums (D) are relatively simple sounds characterized by strong temporal envelope fluctuations and a static spectral structure (harmonic for bass, broadband for drums). These sounds elicit distinct neural events that are coherent across units.

**Figure 1:**
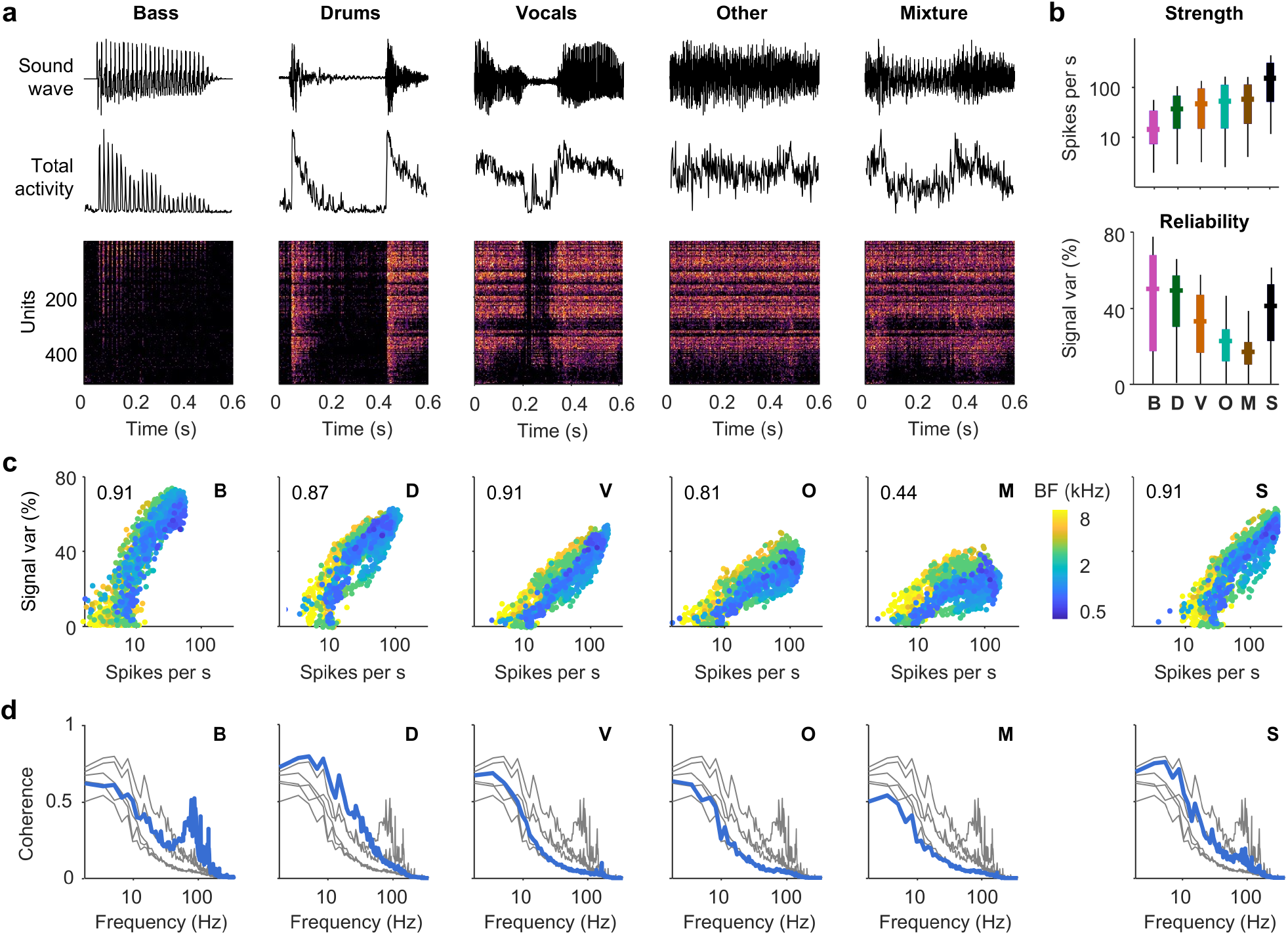
The strength and reliability of neural responses to different music types. **a**. Examples of neural responses to music from an animal with normal hearing. The top row shows the sound waveform, the middle row shows the sum of activity across all units, and the bottom row shows the activity of each unit, with brighter colors indicating more activity and units sorted by preferred frequency (low to high from top to bottom). **b**. The distributions of the strength (top) and reliability (bottom) of neural responses across all units (horizontal line indicates median, thick vertical line indicates 25th through 75th percentile, thin vertical line indicates 5th through 95th percentile; n = 1536). Strength was measured as the overall spike rate. Reliability was measured as the ratio of the signal variance to the total variance (expressed as a percentage), with the signal variance defined as the covariance of responses across trials. Results are shown for each music type (Bass, Drums, Vocals, Other, and Mixture) as well as for the sum of the responses to B, D, V, and O, denoted as S. **c**. The strength and reliability of neural responses for all units, with the dots for each unit colored according to its best frequency (BF; the frequency to which the unit responded most strongly for tones at a range of intensities). **d**. The cross-trial coherence of neural responses, averaged across all units. Each panel shows the coherence spectra for all music types, with the type denoted in the top right of each panel shown in blue and the other types shown in gray.

The vocals (V) are more complex; they also contain strong envelope fluctuations but their spectral structure varies over time from phoneme to phoneme (e.g., harmonic for vowels, broadband for stop consonants) evoking neural activity with a rich structure. (For an extensive analysis of IC responses to speech, see [13, 39].) The other (O) and mixture (M) are the most complex because they are composed of multiple instruments. They are broadband sounds with spectral content that varies over time depending on which instruments are active. Their temporal envelope fluctuations are relatively weak and they elicit persistent neural activity. Thus, the 5 music types can be ordered along a continuum: B, D, V, O, M. The bass and drums are at one end with their strong temporal modulations and simple spectral structure. The other and mixture are at the opposite end with their weak temporal modulations and complex spectral structure. And the vocals lie in between with respect to both temporal modulations and spectral complexity.

We first examined the basic statistical properties of the neural responses to each of the music types for animals with normal hearing (n = 1536 multi-units from 3 animals), assessing how strongly and reliably the units were driven. To assess strength, we measured the overall spike rate. To assess reliability, we measured the percent of the total variance that corresponded to signal rather than noise (i.e., the percent of the variance that was shared across repeated trials of the same music). As shown in Fig. 1b, the strength and reliability of the responses varied inversely across music types, with an ordering that mirrored the continuum of musical complexity established above: the bass responses were the weakest and most reliable, while the mixture responses were the strongest and least reliable.

We also took advantage of the fact that the mixture is the sum of the individual stems (i.e., M = B + D + V + O in the sound domain) to assess the nonlinearity of the integration of the different stems. If auditory processing was linear, the response to the mixture would be the same as the sum of the responses to the individual stems. But auditory processing is, of course, highly nonlinear, and by comparing the true mixture response to the sum of the stem responses (S = B + D + V + O in the neural domain), we can gain insight into how this nonlinearity impacts the neural coding of music. As shown in Fig. 1b, both the strength and the reliability of the mixture responses were much lower than would be expected if the response to the mixture were simply the sum of the responses to the individual stems, indicating a strong impact of nonlinearity on the neural integration of the stems.

When we examined the variation of response strength and reliability across units, we observed several trends (Fig. 1c). The correlation between response strength and reliability was high for the simpler music types, but decreased slightly for the other and decreased substantially for the mixture. The breakdown in the correlation between strength and reliability for the more complex music types appears to result from differential effects across units with different best frequencies (BFs). The correlation between strength and reliability for units with high BFs (green and yellow) appears to be relatively consistent across music types. Units with low BFs (blue), in contrast, are strongly driven for all music types, but their reliability drops sharply for the other and mixture. The breakdown in the correlation between response strength and reliability for the mixture is not evident in the sum of the responses to the individual stems, indicating that it is a result of nonlinear neural integration.

To examine the reliability of the neural responses in more detail, we computed the coherence between responses to repeated trials, which provides a measure of the reliability of each frequency component of the neural responses (Fig. 1d). All of the coherence spectra had higher values for lower frequencies, indicating that slow envelope fluctuations were encoded most reliably, and the variation in the overall magnitude of the coherence across music types reflected the variation in response reliability shown in Fig. 1b. The spectrum for the bass was unique in that it had a second broad peak at higher frequencies corresponding to the fundamental frequencies and low-order harmonics of the instrument. The spectrum for the mixture was similar in shape to the spectrum of the sum of the responses to the individual stems, but was much lower in magnitude. Overall, these initial analyses demonstrate significant variations in neural response properties across music types, revealing clear trends related to musical complexity and the substantial impact of nonlinear neural integration.

### Nonlinear integration distorts the neural coding of music

To gain further insight into the neural integration of the different stems, we computed the coherence between the mixture response and the response to each individual stem (Fig. 2a, step 1). We compared this mix-stem coherence to two benchmarks (Fig. 2a, steps 2 and 3). The first was the cross-trial stem coherence shown in Fig. 1d. If the mix-stem coherence is the same as the cross-trial stem coherence, then the neural coding of the stem is not impacted by the presence of the other stems in the mixture; but if these two coherences are different, then the coding of the stem has been either enhanced or distorted by the other stems.

**Figure 2:**
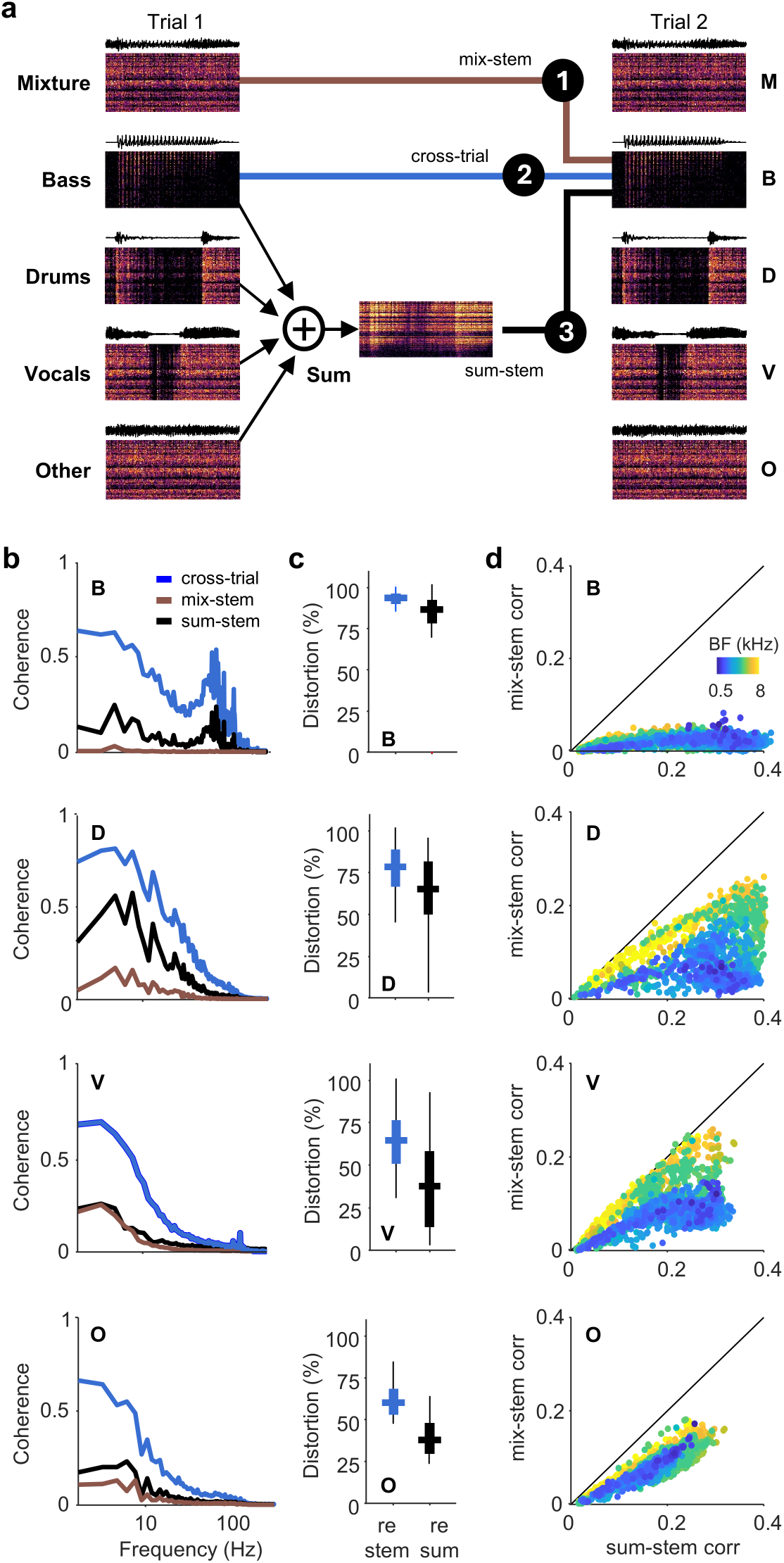
Distortion in the neural coding of mixed music. **a**. A schematic diagram illustrating the three types of comparisons that were made to assess the distortion of the neural representation of individual stems within the response to a mixture: (1) comparison of the response to the mixture with the response to one individual stem (mix-stem); (2) comparison of the responses to an individual stem across repeated trials (cross-trial; the labeled line benchmark); and (3) comparison of the sum of responses to all individual stems with the response to one individual stem (sum-stem; the linear integration benchmark). **b**. The coherence for the comparisons described in a, averaged across all units. **c**. The distributions of distortion of neural responses across all units measured relative to the two benchmarks described in a. Distortion was measured as 1 - the ratio of the correlation between stem and mixture responses to the benchmark correlation (expressed as a percentage). **d**. The correlation between stem and mixture responses versus the linear integration benchmark correlation for all units, with the dots for each unit colored according to its best frequency.

As shown in Fig. 2b, the mix-stem coherence (brown) was much lower than the cross-trial stem coherence (blue) for all stems and all frequencies, indicating a high degree of distortion. As a summary metric, we computed the ratio of the correlation between the stem and mixture responses to the correlation between the stem responses across trials, and then subtracted the ratio from 1 to obtain a measure of the total distortion (expressed as a percentage). By this metric, distortion decreased with increasing complexity, i.e., from bass to other; the distortion of the neural representation of the bass within the mixture response was near complete, while for the other it was approximately 60% (Fig. 2c).

The use of the cross-trial stem coherence as a benchmark reflects an ideal scenario in which the different stems of the mixture are encoded by distinct neural populations that do not interact (i.e., labeled lines), which is unlikely given the overlapping spectral content of the different stems. As a second benchmark that reflects more reasonable expectations, we used the coherence between the mixture response and the sum of the responses to the individual stems. If the mix-stem coherence is the same as the sum-stem coherence, then the distortion of the neural representation of the stem within the mixture response is equivalent to that expected for linear neural integration; if these two coherences are different, then the coding of the stem has been nonlinearly enhanced or distorted by the other stems in the mixture.

Comparison against this benchmark was more favorable, but the total distortion (measured as described above) was still at least 40% for all stems (Figs. 2b,c). Distortion again decreased with increasing stem complexity, with the bass experiencing the most distortion and the other experiencing the least. When examining the variation in distortion across units, we again observed several trends (Fig. 2d). The distortion exhibited by all units decreased with increasing musical complexity, but this decrease was more rapid for units with high BFs, i.e., for the drums and vocals, the distortion exhibited by units with high BFs was lower than that exhibited by units with low BFs. Taken together, the results thus far demonstrate that the combination of stems creates interactions that are highly nonlinear and detrimental to the neural representation of the individual stems, with particularly strong effects for the simpler stems and units with low preferred frequencies.

### Increasing musical complexity creates new neural modes

The coherence analyses described above treat each neural unit as an independent information channel. A full characterization of neural coding, however, requires direct analysis of activity patterns at the network level. One simple but important aspect of the neural code that is ignored in the coherence analyses is the variation in overall spike rate across units, which will, of course, reflect the different acoustic properties of the different music types. For the simpler music types (bass and drums) with their strong temporal envelope fluctuations that are coherent across frequencies, we might expect relatively strong variations in spike rates over time but relatively weak variation in overall spike rate across units. For the more complex music types (other and mixture), with their varied spectral content and relatively weak temporal envelope fluctuations, we might expect the opposite. When we computed the fraction of the signal variance in the network-level activity patterns that was accounted for by the variation in overall spike rate across units, we found that these expectations were met (Fig 3a; a value of 50% would indicate that the reliable variations in overall spike rate across units are as strong as the reliable variations in the spike rates of each unit over time). This fraction was lowest for the bass and highest for the mixture, with an ordering that again mirrored the B, D, V, O, M continuum of musical complexity.

**Figure 3:**
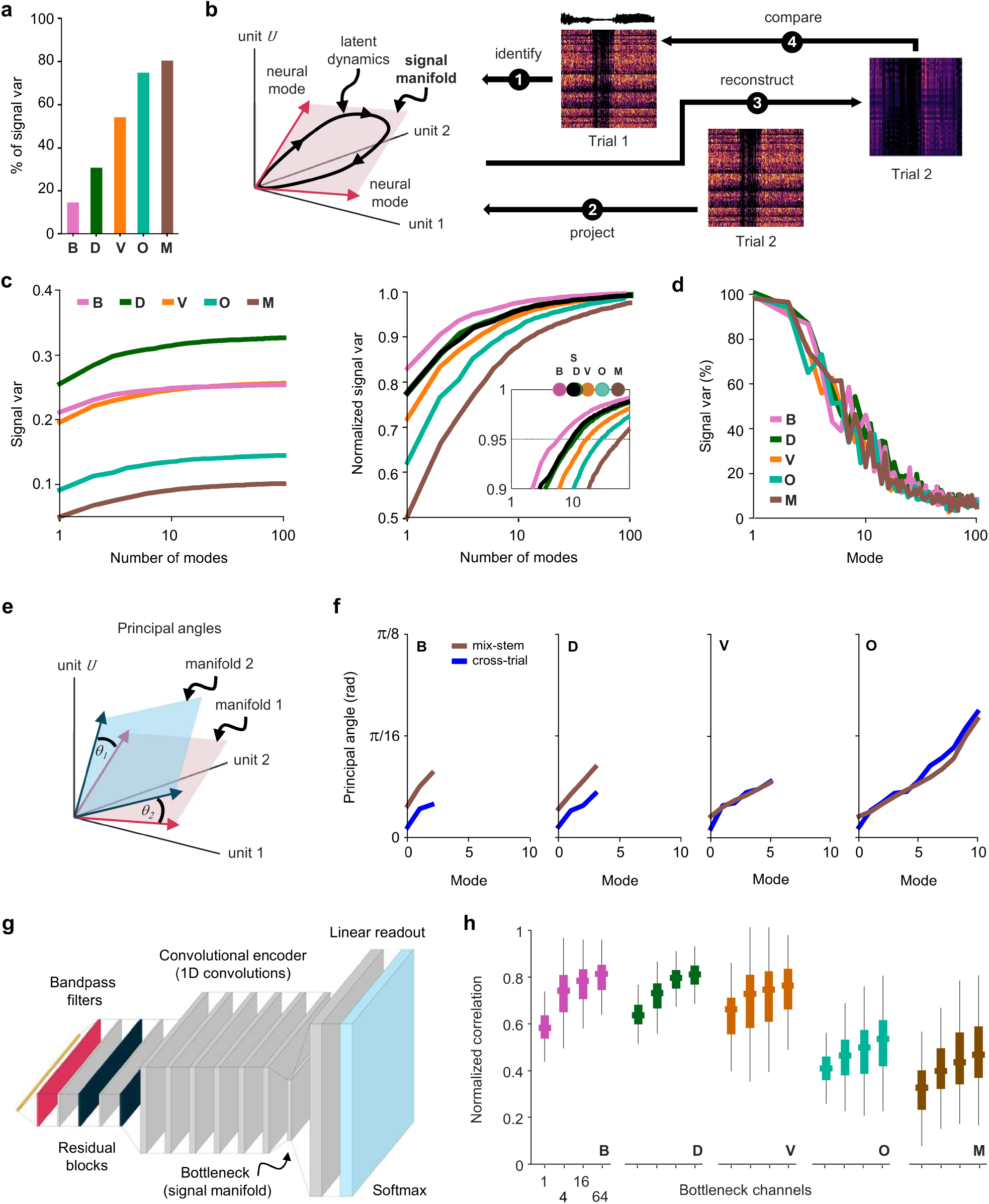
Manifold analysis and modeling of the neural code for music. **a**. The percentage of the signal variance in the full responses across all units that is accounted for by the variation in overall spike rate across units. **b**. A schematic diagram illustrating the identification and analysis of the signal manifold. PCA was performed on responses to one trial; the responses for a second trial were projected onto a subset of the identified components (modes) and then reconstructed in the full response space; and the covariance between the reconstructed responses and the full responses from the first trial was computed. The signal manifold was defined as the subspace spanned by the modes that captured 95% of the full signal variance, i.e., the covariance between the full responses across trials. **c**. The covariance between the full responses from one trial and the reconstructed responses from another trial as a function of the number of neural modes used. The left panel shows the raw values and the right panel shows the values after normalization by the covariance between the full responses across trials. The inset in the right panel shows dots indicating the number of modes required to captured 95% of the full signal variance for each music type as well as for the sum of the responses to the stems. **d**. The percent of the variance captured by each neural mode that corresponds to neural signal rather than neural noise. **e**. A schematic diagram illustrating the use of principle angles to measure the alignment of neural subspaces. **f**. The principal angles between the signal manifolds for the mixture and each stem (brown), along with the principal angles between the signal manifolds identified for each stem on separate trials (blue). All angles were computed for the subspaces spanned by the first 15 modes, but for each music type only the modes required to capture 90% of the full signal variance are shown. **g**. The architecture of the deep neural network (DNN) that was trained to predict neural responses to music. An encoder-decoder model was trained in a classification framework to predict the probability distributions of possible spike counts for each time bin and unit using cross-entropy loss. Responses were simulated from the trained model by sampling from the inferred distributions. **h**. The performance of the DNN models as a function of the number of channels in the bottleneck for each of the music types. Performance was measured as normalized correlation, the ratio of the correlation between the model prediction and the actual responses to the correlation between the actual responses on repeated trials.

To examine the network-level activity patterns in detail, we utilized manifold analysis [7, 11, 31] (Fig. 3b). We first identified the signal manifold (the low-dimensional subspace in which the reliable features of neural activity are embedded) for each music type using principal component analysis (PCA). To focus on features of the neural code beyond overall spike rate, we centered the neural activity for each unit before applying PCA. In the IC, the noise in neural activity is independent across units [13, 39], while the signal may be shared across units if their acoustic preferences are similar. Since PCA identifies components that capture variance that is shared across variables, it preferentially identifies neural modes in IC activity that are associated with signal rather than noise [39].

After identifying the signal manifold from responses to one trial of a specific music type, we projected the responses from a second trial of the same music onto this manifold and measured how much of the first trial’s variance was captured by the projection (of course, only signal variance can be captured since the noise in the responses will be independent across trials). To determine the dimensionality of each signal manifold, we measured the number of neural modes, i.e., principal components (PCs), required to capture 95% of the full signal variance (Fig. 3c). This dimensionality varied widely across music types and increased with increasing musical complexity, with a dimensionality of 6 for the bass and 52 for the mixture. And the dimensionality of the signal manifold for the sum of the responses to the individual stems (10) was much lower than that for the mixture.

These results suggest that the nonlinearity of auditory processing drives the creation of new neural modes as musical complexity is increased. But it is also possible that the apparent high dimensionality of the manifolds for the more complex music types is a simple consequence of the relatively low reliability of the responses to these types (Fig. 1b). While PCA of IC activity preferentially identifies neural modes that carry signal rather than noise, the projection of neural responses onto those modes can never separate signal from noise completely. If the leading modes (i.e., the top PCs) of the manifolds for the more complex music types are noisier than those of the manifolds for the simpler music types, then more modes would be required to capture a given fraction of the signal variance. However, when we measured the fraction of the variance associated with each neural mode that corresponded to signal rather than noise, we found that the values were nearly identical for all music types (Fig. 3d).

To gain further insight into how musical complexity impacts the geometry of neural representations, we measured the principal angles [12] between the leading modes of the identified signal manifolds (Fig. 3e). (These angles measure the alignment between the subspaces spanned by the modes; principal angles close to zero indicate that the subspaces are closely aligned, while principal angles close to *π*/2 indicate that the subspaces are nearly orthogonal.) As shown in Fig. 3f, the principal angles between the manifolds of the individual stems and the manifold of the mixture were small, and comparable to the angles observed between the manifolds for the same stems estimated from responses to different trials. (Another measure, the correlations of the weights associated with individual units for the leading modes, also indicated high similarity across music types; see Fig. S1). Inasmuch as there are differences between the manifolds for the individual stems and the manifold for the mixture, they appear to be largest for the bass and drums, consistent with the fact that these stems exhibit the highest level of distortion in the mixture response (Fig. 2b-d).

Taken together, the results of the manifold analysis suggest that the leading modes of the subspace in which music is encoded are relatively stable across music types. But with increasing musical complexity, additional modes are created and variance is displaced from the leading modes onto these new modes. Because the new neural modes recruited to encode complex music are less reliable than the leading modes, this increase in dimensionality comes at the expense of reduced overall response reliability..

### Deep neural network models capture the tradeoff between reliability and dimensionality

As a further step toward characterizing the neural signal manifold for each music type, we modeled the mapping from music to neural activity using a deep neural network (DNN). The model featured an encoder-decoder architecture with a low-dimensional bottleneck (Fig. 3g). The encoder comprised a series of convolutional layers, allowing a nonlinear mapping from the input sound to the latent representation. The decoder, in contrast, used a simple linear readout to maintain a transparent link between the latent representation and the output neural activity [39, 8]. We trained separate models for each music type and evaluated the models on music of the same type while assessing the dependence of the model performance on the number of channels in the latent bottleneck.

The models performed well overall, with the average normalized correlation (the correlation between the model response and the actual response divided by the correlation between the actual responses across trials) ranging from 0.48 to 0.8 depending on music type for the models with the largest bottlenecks (Fig. 3h; see Fig. S2 for examples). If the latent representations learned by the DNNs are comparable to those identified via PCA, then we should expect the dependence of the model performance on the number of bottleneck channels to reflect the dimensionalities of the signal manifolds reported above, and this was indeed the case. For the bass, which had the lowest signal manifold dimensionality, the model performance improved markedly when increasing from 1 to 4 bottleneck channels but only gradually thereafter, whereas for the mixture, which had the highest signal manifold dimensionality, the performance increased gradually across the full range of bottleneck channels.

It is also evident from Fig. 3h that even with the largest number of bottleneck channels (64, which exceeds the dimensionality of any of the signal manifolds), the model performance varied widely across music types with an ordering that followed the B, D, V, O, M continuum of musical complexity. That the ordering of model performance across music types matched that of the reliability of neural responses rather than that of the response strength (Fig. 1b) suggests that the former rather than the latter is the key determinant of how readily a mapping from sensory stimulus to neural response can be learned. We also trained a master model on all music types, which performed similarly to the type-specific models for all music types and exhibited the same B, D, V, O, M ordering of performance across types (Fig. S2). This is consistent with the idea that the signal manifolds for the different music types are similar (e.g., that the leading modes are largely shared). Both Python and Matlab versions of this master model will be made freely available upon publication to allow other researchers to use it for applications that can benefit from brain-inspired representations of music.

### The effects of hearing loss on the neural coding of music are largely corrected by amplification

To gain insight into how hearing loss changes the neural coding of music, we recorded and analyzed responses to the same music from gerbils that were exposed to broadband noise (n = 1536 multi-units from 3 animals). The noise exposure resulted in hearing loss that was mild-to-moderate with threshold shifts of 30-50 dB for frequencies from 1-8 kHz (Fig. 4a). If music is presented to a listener with hearing loss at a sound level that is comfortable for a normal hearing listener, some of the effects of hearing loss will be related to decreased audibility. While these effects are important, they are also easily addressed through amplification. To decouple the effects of decreased audibility from the other effects of hearing loss, we compared the activity from normal hearing (NH) animals with that from animals with hearing loss (HL) at either equal sound level (74 dB SPL for the mixture) or at a higher sound level at which the strength of the neural responses was restored to normal. To choose this higher level, we presented the music at a range of intensities to find the intensity at which the overall HL spike rate was closest to the overall NH spike rate at 74 dB SPL. We settled on 98 dB SPL, which was the lowest level at which the overall HL spike rate exceeded the overall NH spike rate (Fig. 4b).

**Figure 4:**
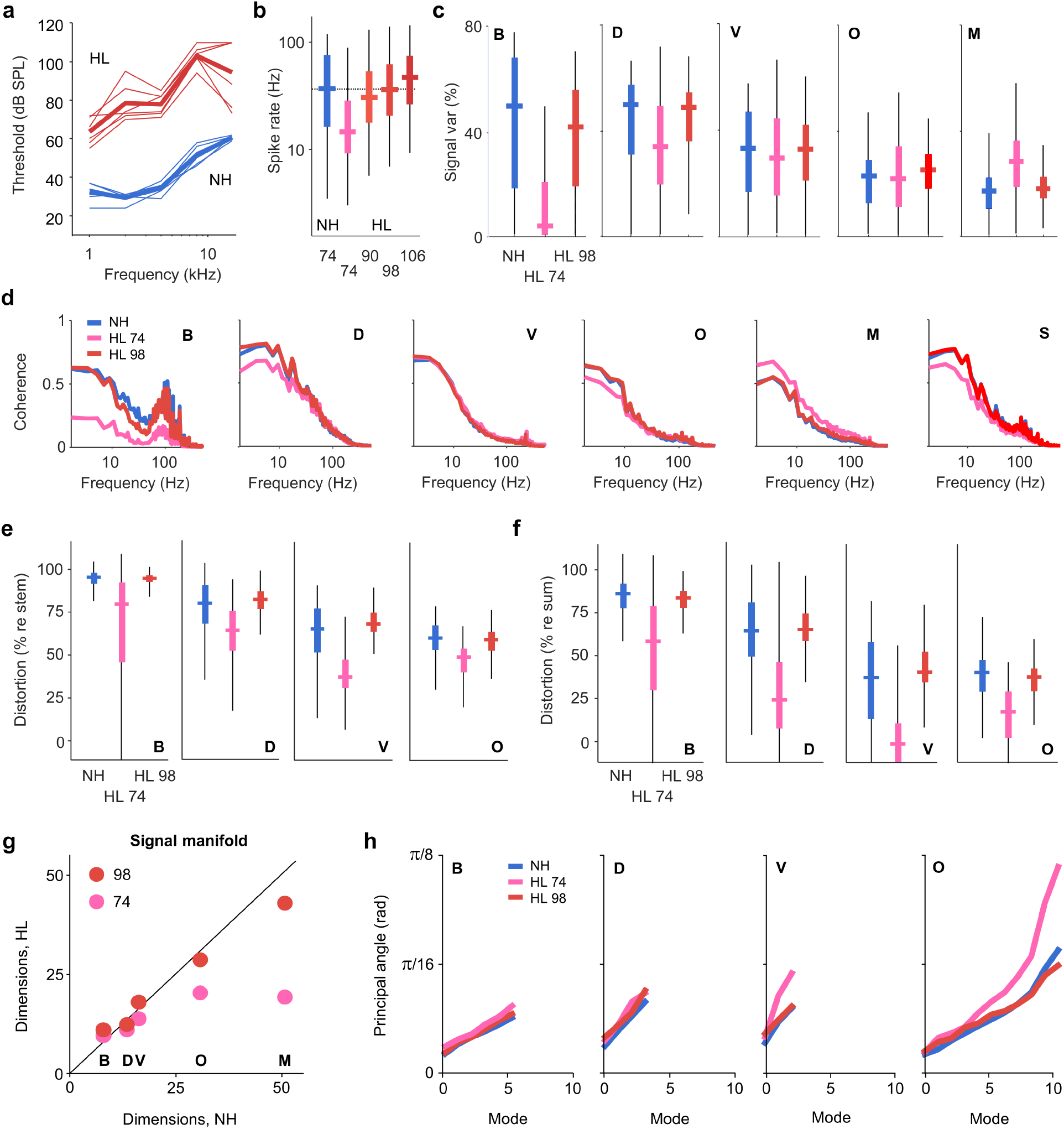
The effects of hearing loss on the neural coding of music. **a**. ABR threshold for ears from gerbils with normal hearing (blue) and hearing loss (red). The thin lines are individual ears and the thick lines are the average across ears. **b**. The distributions of response strength, measured as overall spike rate averaged across all music, for units from normal hearing gerbils (n = 1536) at 74 dB SPL and units from gerbils with hearing loss (n = 1536) at varying intensities. **c**. The distributions of response reliability for units from normal hearing gerbils at 74 dB SPL (blue) and units from gerbils with hearing loss at 74 dB SPL (pink) and 98 dB SPL (red). **d**. The cross-trial coherence of neural responses, averaged across all units, from normal hearing gerbils at 74 dB SPL and from gerbils with hearing loss at 74 dB SPL and 98 dB SPL. **e**,**f**. The distributions of distortion of neural responses across all units measured relative to the two benchmarks described in Fig. 2. from normal hearing gerbils at 74 dB SPL and from gerbils with hearing loss at 74 dB SPL and 98 dB SPL. **g**. The number of modes required to captured 95% of the full signal variance for each music type. The values for normal hearing gerbils at 74 dB SPL are plotted against those for gerbils with hearing loss at 74 dB SPL and 98 dB SPL. **h**. The principal angles between the signal manifolds for the mixture and each stem for normal hearing gerbils at 74 dB SPL and for gerbils with hearing loss at 74 dB SPL and 98 dB SPL.

At the level of units, hearing loss had strong effects on neural coding but these effects were almost entirely related to audibility. At 74 dB SPL, hearing loss impacted the reliability of responses to the bass (reduced relative to normal) and the mixture (increased relative to normal) but these differences, as well as the smaller differences for the other music types, were largely eliminated when the level of the music was increased to 98 dB SPL (Fig. 4c). The variations in overall spike rate and reliability across units with hearing loss were also similar to those observed with normal hearing, albeit with an absence of units with very high BFs (Fig. S3), and the average cross-trial coherence spectra with and without hearing loss were nearly identical when compared at matched overall spike rates (Fig. 4d).

Comparisons of neural distortion with and without hearing loss showed similar trends. There were again differences when the music was presented at 74 dB SPL, but when the sound level was increased, these differences largely disappeared (Figs. 4e,f). The variations in the level of distortion across units with hearing loss were also similar to those observed with normal hearing, with units with lower BFs exhibiting more distortion relative to that expected for linear integration than units with higher BFs (Fig. S3). Thus, it appears that the effects of hearing loss on neural coding at the level of individual units are largely eliminated with suitable amplification.

Audibility also accounted for most of the effects of hearing loss on neural coding at the network level. The dimensionality of the signal manifolds was lower with hearing loss when the music was presented at 74 dB SPL, at least for the more complex music types, but when the sound level was increased, the dimensionality of the signal manifolds returned to normal (Fig. 4g). As shown in Fig. 4h, the principal angles between the leading modes of the signal manifolds for the mixture and the individual stems were nearly identical with and without hearing loss, with only minor differences across sound levels with hearing loss, indicating that the leading modes of the neural subspace in which music is encoded are remarkably stable. Thus, it appears that the effects of hearing loss on the neural coding of music are due almost entirely to the loss of audibility, at least for the mild-to-moderate level of hearing loss that we induced.

## Discussion

Our results provide the first comprehensive characterization of the neural coding of music. By using different music types with a range of spectrotemporal properties, including a mixture that was the sum of stems that were presented individually, we found that the neural code for music is shaped by nonlinear interactions at both the unit and network level. For music consisting of single instruments with strong, spectrally coherent envelope fluctuations and a stable spectral structure (bass and drums), neural responses were weak but reliable, and the shared variance across units could be captured by a small number of neural modes. For music consisting of multiple instruments with weak envelope fluctuations and a varying spectral structure (other and mixture), neural responses were stronger but less reliable, and the shared variance was spread across many more neural modes. For vocals, which have strong envelope fluctuations and a varying spectral structure, the neural responses sat midway between these extremes.

These results suggest that a tradeoff between reliability and dimensionality that is governed by spectrotemporal complexity is a fundamental aspect of the neural code for music. It is important to recognize, however, that our neural recordings from the IC of anesthetized animals cannot capture the influence of certain factors beyond acoustics that may be important for the neural coding (and perception) of music in humans, such as active or contextual processing or the impact of musical training or preference. But the gerbil and human early auditory pathways are similar in many ways that are relevant for neural coding, and the influence of top-down processes and experience on peripheral processing of music appear to be relatively modest [10, 46]. So it is reasonable to assume that the fundamental tradeoff that we observed would also be present in the neural code in the IC of humans, albeit with potential modulation by other factors.

### Linking the neural coding of music to perception

We found that the neural representation of an individual stem is profoundly altered by the presence of other stems within a mixture. Inasmuch as the neural representation in the early auditory pathway establishes the basis for perception, our results can be viewed as possible neural correlates of phenomena that have been observed in studies of musical scene analysis. For example, when listeners were asked to detect individual instruments within a mixture of bass, guitar, piano, and vocals, the bass proved to be the most difficult to detect [15]. This is consistent with our finding that the neural representation of the bass was the most distorted of all the stems when presented in a mixture.

Strong interactions between stems in the neural coding of music are to be expected; there must be some complex aspects of auditory processing that help to make mixed music so pleasant to listen to and so challenging to compose. But the low reliability of the neural responses to mixed music is surprising. While responses to the mixture were strongly driven, only a small fraction of their variance was repeatable across presentations. Yet, somehow, these noisy neural responses give rise to a perceptual experience of high fidelity. This suggests that there are high-level processes that are effective at separating neural signal from neural noise or creating invariance to neural noise in some other way.

Identifying the neural mechanisms responsible for converting noisy neural responses into a high-fidelity percept would be a key step toward understanding the link between neural coding and perception. The results of our manifold analysis provide one possible direction to explore. We found that the dimensionality of the neural signal manifold increased with musical complexity, with the mixture exciting many more neural modes than any of the individual elements. But we also found that the leading modes, and the fraction of neural signal and noise associated with them, remained stable across music types. This suggests that downstream processing that is focused on the leading modes could be invariant to the additional noise associated with higher modes. But such processing would, of course, also miss out on the signal carried by the higher modes. Further exploration of the impact of the dimensionality of neural representations on perception could provide some clarity. For example, experiments could assess how well listeners can distinguish music reconstructed from responses projected onto an increasing number of neural modes. The results of such experiments could reveal whether the signal carried by higher modes is perceptually relevant.

### The effects of hearing loss on the neural coding and perception of music

We found that effects of hearing loss on the neural code for music, whether at the unit or network level, were almost entirely eliminated by providing appropriate amplification. When presenting music to animals with normal hearing and hearing loss at the same sound level (74 dB SPL), we observed large differences in the strength, reliability, distortion, and dimensionality of neural responses. But after increasing the sound level to 98 dB SPL so that the responses with hearing loss were of normal strength, we found that the abnormalities in the other response properties were also largely corrected. Interestingly, the principal angles between the neural subspaces defined by the leading modes of signal manifolds for the mixture and the individual stems were similar across all of the conditions that we tested, suggesting that the core geometry of the neural code is remarkably stable. While our comparisons of the neural code with and without hearing loss are indirect, i.e., we made comparisons across different animals rather than before and after hearing loss in the same animals, our results suggest that the effects of hearing loss on the neural coding of music are largely related to audibility.

The limited effects of hearing loss on neural coding may seem surprising given the widely documented effects of hearing loss on music perception [32, 2, 27]. However, it may be that the severity of the hearing loss that we induced – mild-to-moderate with threshold shifts of 30-50 dB – is not sufficient to elicit complex effects on the neural coding of music. This level of hearing loss is not trivial, and is associated with deficits in the neural coding and perception of speech-in-noise [9, 18, 28]. But for the neural coding and perception of music, it may be that more severe losses are required to elicit effects beyond those related to audibility. For example, in the same study of musical scene analysis described above, the performance of listeners with hearing loss (who were able to self-adjust the sound level) was only significantly worse than normal when the level of hearing loss was beyond moderate, i.e., severe or profound [15]. In light of these results, current hearing aid music programs, which are relatively simple amplification schemes [5], seem well matched to the needs of listeners with relatively mild loss. Further work is required to understand the impact of more severe hearing loss on the neural coding of music and to determine what additional processing beyond amplification might be beneficial.

### Relationships between complexity, reliability, and dimensionality in neural systems

One starting point for understanding the observed variation in neural responses across music types is to consider two basic acoustic features: the modulation depth of envelope fluctuations and the correlation across spectral bands. For the bass and drums, these features are strong; for the other and mix, they are weak. Even at the most peripheral level of the auditory system, the reliability of neural responses is strongly dependent on envelope modulation depth (i.e., the rate synchronization of auditory nerve (AN) fibers increases with increasing modulation depth [20]). And while the dimensionality of AN responses has not been explicitly studied, it is likely that higher correlation across spectral bands leads to more coordinated activity along the cochlea and, consequently, more shared variance among the population of AN fibers that can captured by fewer neural modes. Thus, the tradeoff that we observed in IC responses is likely already established in the cochlea based on these basic acoustic features. There may also be contributions from other factors, either higher-order acoustic properties or neural mechanisms within the brain itself. Further studies of neural coding at different processing stages, using artificial sounds in which envelope fluctuations and spectral correlations can be decoupled from each other and from other acoustic properties, could establish the degree to which simple acoustic features and peripheral processing are sufficient to explain the observed tradeoff.

It is also possible that the relationship between music complexity and the reliability and dimensionality of neural responses reflects more general constraints on neural coding. This would be consistent with basic concepts in systems theory and complexity science suggesting that a system’s behavior is more stable when its constituent elements are able to work in concert. Related phenomena have also been observed in other sensory systems. For example, the reliability of neural responses in the early visual pathway varied across stimulus types, with the highest reliability for high-contrast full-field flicker (with strong spatially-coherent fluctuations) and the lowest reliability for low-contrast checkerboard patterns (with weak spatially-incoherent fluctuations) [6].

Another intriguing possibility is that the constraints that link input complexity, latent dimensionality, and output reliability extend beyond natural neural systems to artificial neural networks. That the properties of a DNN would accurately reflect the properties of the data on which it is trained is to be expected. But we also found that the overall performance of our model (based on a metric that normalized for neural response reliability) varied across music types with an ordering that mirrored that for response reliability. This may be a simple consequence of the signal-to-noise ratio in the data (the mapping from sound to response may be harder to learn from noisier responses). But it may also reflect something more subtle about the degree to which a mapping of a given complexity can be learned by a model with a given capacity. Modeling experiments in which the complexity, dimensionality, and signal-to-noise ratio of input-output mappings are systematically manipulated could reveal important connections between artificial and biological networks.

## Methods

### Experimental protocol

Experiments were performed on 6 young-adult gerbils (3 female, 3 male) that were born and raised in standard laboratory conditions. Three of the animals were exposed to noise when they were 16-18 weeks old. ABR recordings and large-scale IC recordings were made from all animals when they were 20-24 weeks old. All experimental protocols were approved by the UK Home Office (PPL P56840C21). IC recordings were analyzed at the unit and network level using coherence analysis, manifold analysis, and deep neural network modeling as described below.

### Noise exposure

Mild-to-moderate sensorineural hearing loss was induced by exposing anesthetized gerbils to high-pass filtered noise with a 3 dB/octave roll-off below 2 kHz at 118 dB SPL for 3 hr [1, 43]. For anesthesia, an initial injection of 0.2 ml per 100 g body weight was given with fentanyl (0.05 mg per ml), medetomidine (1 mg per ml), and midazolam (5 mg per ml) in a ratio of 4:1:10. A supplemental injection of approximately 1/3 of the initial dose was given after 90 min. Internal temperature was monitored and maintained at 38.7°C.

### Preparation for large-scale IC recordings

Animals were placed in a sound-attenuated chamber and anesthetized for surgery with an initial injection of 1 ml per 100 g body weight of ketamine (100 mg per ml), xylazine (20 mg per ml), and saline in a ratio of 5:1:19. The same solution was infused continuously during recording at a rate of approximately 2.2 *µ*l per min. Internal temperature was monitored and maintained at 38.7°C. A small metal rod was mounted on the skull and used to secure the head of the animal in a stereotaxic device. The pinnae were removed and speakers coupled to tubes were inserted into both ear canals along with microphones (Etymotic ER-10X) that were used to calibrate the sound delivery. The frequency response of the sound system measured at the entrance of the ear canal was flat (±5 dB) between 0.2 and 8 kHz. Two craniotomies were made along with incisions in the dura mater, and a 256-channel multi-electrode array was inserted into the central nucleus of the IC in each hemisphere [1, 39].

### Auditory brainstem responses

Before beginning the IC recordings, ABRs were measured. Subdermal needles were used as electrodes with the active electrodes placed behind the ear over the bulla (one on each side), the reference placed over the nose, and the ground placed in a rear leg. Recordings were bandpass filtered between 300 and 3000 Hz. The parallel ABR method was used [35, 44], with randomly timed tones at multiple frequencies presented simultaneously and independently to each ear. The tone frequencies were 1, 2, 4, 8, and 16 kHz. Each tone was 5 cycles long and multiplied by a Blackman window of the same duration. Tones were presented at a rate of 40 per s per frequency with alternating polarity for 100 s at each intensity. The activity recorded in the 30 ms following each tone was extracted and thresholds for each frequency were defined as the lowest intensity at which the root mean square (RMS) of the median response across presentations was more than twice the RMS of the median activity recorded in the absence of sound.

### Auditory stimuli

#### Tones

We presented 50 ms tones with frequencies ranging from 300 Hz to 16000 Hz in 0.2 octave steps and intensities ranging from 4 dB SPL to 85 dB SPL in 9 dB steps with 2 ms cosine on and off ramps and 175 ms pause between tones. Tones were presented 8 times each in random order.

#### Music

We presented music from *musdb18*, a freely-available benchmark dataset designed for audio source separation research [36]. It consists of 150 full-length music tracks of various genres, each professionally recorded and manually mixed. Each track is provided in stem format, meaning it is separated into four distinct components or “stems”: bass, drums, vocals, and other (everything else). The dataset is split into a training set of 100 songs and a test set of 50 songs. For normal hearing animals, we presented each of the stems and the mixture separately for 23 full pop/rock songs from the training set (8.3 hr) one time each, as well as the stems and mixture for 15 seconds from 25 pop/rock songs from the test set (31.3 min) two times each, all at 74 dB SPL. For animals with hearing loss, we presented only the music from the test set but at range of different intensities.

### Analysis of recorded neural activity

#### Multi-unit activity

MUA was extracted from recordings on each channel of the electrode array as follows: (1) a bandpass filter was applied with cutoff frequencies of 700 and 5000 Hz; (2) the standard deviation of the background noise in the bandpass-filtered signal was estimated as the median absolute deviation / 0.6745 (this estimate is more robust to outlier values, e.g., neural spikes, than direct calculation); (3) times at which the bandpass filtered signal made a positive crossing of a threshold of 3.5 standard deviations were identified and grouped into bins with a width of 1.3 ms.

All analyses aside from DNN training were carried out on the responses to the two presentations of the music from the test set (see ‘Music’ above). Some individual stems included extended periods of silence. The data were divided into non-overlapping 7.5 s segments and the segments that were completely silent for each stem were identified. For the analyses in which responses to different music types were treated seperately (Figs. 1 and 4a-d), all non-silent segments were included for each stem, meaning the number of time bins could differ across the stems and the mixture. For the analyses in which the responses to different music types were directly compared to each other (Figs. 2, 3, and 4e-h), only those segments that were non-silent across all stems were used.

We denote the neural response matrices as *R*_*xy*_, where *x* is the music type (*b* for bass; *d* for drums; *v* for vocals; *o* for other; *m* for mixture; and *s* for the sum of the neural responses to *v, d, b* and *o*) and *y* is the trial (1 or 2). Each *R*_*xy*_ is a *U × T* matrix, where *U* is the number of units (1536 for each hearing condition), *T* is the number of time bins, and each row 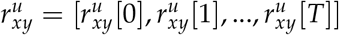 is the vector of spike counts for unit *u*.

#### Strength and reliability

The basic statistical properties of the neural responses to each music type were evaluated by quantifying the overall strength and reliability of unit activity.The overall strength was assessed by computing the mean spike rate for each unit, defined as

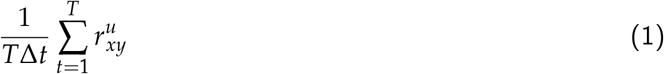

The reliability was assessed by computing the percentage of the variance in the neural response that was repeatable across trials, defined as

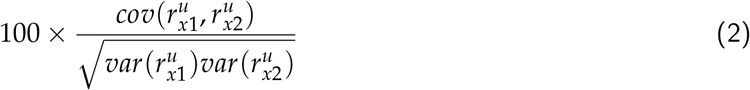

The reliability of the individual frequency components of the neural responses was also assessed using the magnitude-squared coherence between responses across trials. The coherence 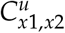 was defined as

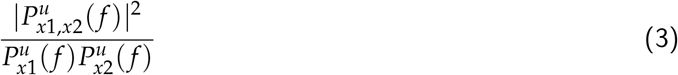

where cross spectral densities 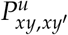 of 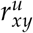 and 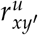 and power spectral densities 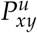 of 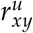 were computed using Welch’s averaged periodogram method with 1024 samples for each discrete Fourier transform, a Hamming window with 2048 samples, and an overlap of 1024 samples between segments.

#### Distortion

To measure the extent to which the neural coding of individual stems was enhanced or distorted by the other stems in the mixture, we compared the responses to the individual stems with the responses to the mixture or the sum of the responses to the individual stems. We computed the coherence as described above to obtain either the sum-stem coherence 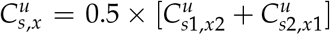 or the mix-stem coherence 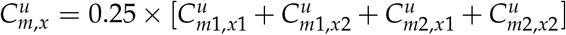. We also computed the overall distortion relative to the sum-stem benchmark as

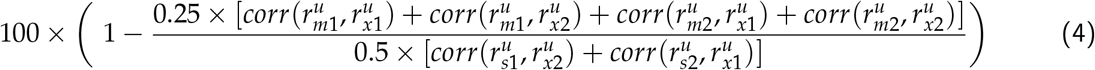

and the overall distortion relative to the cross-trial benchmark as

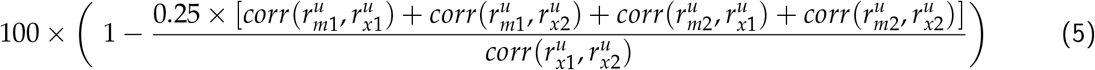

#### Signal variance accounted for by overall spike rates

To determine how much of the signal variance in the full time-varying neural responses across units could be accounted for by the differences in the overall spike rates across units we computed

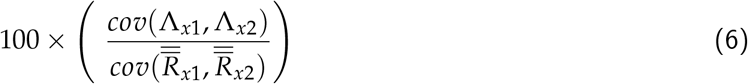

where 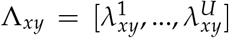 is the vector of mean spike counts for each unit 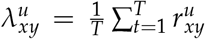 and the double bar operator denotes flattening, i.e., the reshaping of a matrix into a column vector.

#### Identification of the signal manifold

We applied PCA to the response to each trial a given music type *R*_*xy*_, resulting in *W*_*xy*_, a *U × U* weight matrix of principal components (referred to as neural modes) with each row containing the weights for one mode. The neural activity for each unit was centered before PCA was applied. We projected the activity *R*_*xy*_ onto the first *d* modes of 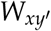 to obtain the latent representation of the neural dynamics from trial *y* in the signal manifold estimated from trial 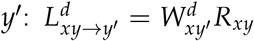 where 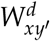 contains the first *d* rows of 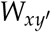. We reconstructed full neural activity from the projection as 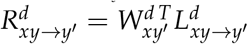.

We measured the signal variance captured by a set of modes as 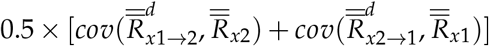 with *d* varied from 1 to *U*. We defined the dimensionality of the signal manifold as the lowest value of *d* for which the signal variance exceeded 95% of the signal variance in the full responses 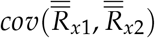.

#### Reliability of neural modes

We measured the reliability of the latent dynamics associated with each neural mode as

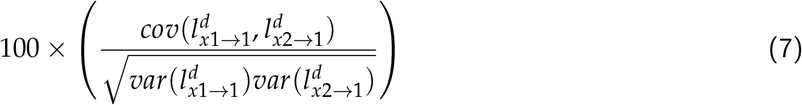

where 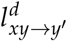 is the *d*^*th*^ row of 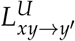.

#### Geometry of the signal manifold

To assess the similarity of the signal manifolds for the different music types, we used principal angles as a measure of alignment between subspaces. We computed the inverse cosines of the singular values of the outer product of two sets of modes, using only the first 15 modes. The mix-stem principal angles were computed as *θ*_*m,x*_ = 0.25 *× θ*_*m*1,*x*1_ + *θ*_*m*1,*x*2_ + *θ*_*m*2,*x*1_ + *θ*_*m*2,*x*2_ where 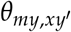 is the vector of the inverse cosines of the singular values of 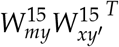.

### Deep neural network models

#### Model architecture

We developed a DNN to predict neural activity from sound waveforms. Following previous efforts to model neural coding in the IC [39, 8], we used a fully convolutional encoder-decoder architecture to map time domain sound waveforms to MUA spike count probabilities. The model takes as input raw audio with shape (8192, 1), corresponding to ~335 ms at a sampling rate of 24,414.0625 Hz. The model architecture comprised the following:

- A SincNet layer, which applies a bank of 48 parameterized sinc filters with a kernel size of 32, designed to mimic the band-pass filtering as observed in the cochlea.
- Logarithmic amplitude compression, which applies a variant of symmetric log activation to the filtered signals *x* = *sign*(*x*)*log*(1 + |*x*| + *ϵ*) with *ϵ* = 1*x*10^−6^ to ensure numerical stability.
- A stack of two residual blocks, each consisting of a 1-D convolutional layer with 48 filters and a kernel size of 32 followed by PReLU activation, with the residual summation made before the PReLU.
- A 1-D convolutional layer with 128 filters, a dilation rate of 2, and ELU activation.
- A stack of five 1-D convolutional layers, each with 128 filters, a kernel of size 32, a stride of 2, and PReLU activation.
- A bottleneck 1-D convolutional layer with a variable number of filters, a kernel of size 32, a stride of 1, and PReLU activation.
- A decoder 1-D convolutional layer with 1536 × 5 filters (one for each unit and possible spike counts from 0 to 4), a kernel size of 1, no bias, and softmax activation across spike counts.

The model produces as output spike count probabilities with shape (256, 1536, 5), which was cropped to 192 samples along the first dimension to remove convolutional edge effects.

#### Model training and evaluation

Models were trained on the neural responses to the music from the training set (see ‘Music’ above) with 15% of the data reserved for validation after each epoch. Models were trained on NVIDIA RTX 3080 GPUs using a batch size of 64 with the Adam optimizer and a learning rate of 0.0004. Training was carried out until the validation loss failed to decrease for 4 consecutive epochs or a maximum of 30 epochs was reached.

To evaluate performance, we used the model to process music from the test set and sampled from the inferred spike count distributions to generate MUA. We assessed performance by computing the correlation between model-generated activity and the neural responses. To account for the cross-trial variability in the responses (i.e., neural noise), we normalized this correlation by the correlation of the neural responses across repeated trials. This normalized measure assesses the performance of the model relative to the ceiling imposed by the noise in the data.

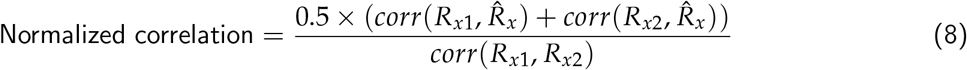

where 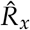 is the model-generated MUA response to sound *x*.

## Acknowledgements

This work was supported by a grant from the UK Medical Research Council (MR/W019787/1). The authors thank A. de Cheveigné, Gerardo Roa Dabike, Michael Akeroyd, Jennifer Firth, Scott Bannister, and Trevor Cox for advice and preparation of music materials.

## Author contributions

P.G.M.: Formal analysis, Methodology, Software, Validation, Visualization, Writing: Original Draft Preparation; S.S.: Investigation; A.E.G.: Conceptualization, Funding acquisition, Project administration, Supervision, Writing-Review & Editing; N.A.L.: Conceptualization, Data curation, Funding acquisition, Investigation, Methodology, Project Administration, Supervision, Writing: Original Draft Preparation

## Data availability

The IC recordings that were used in this study are available on Zenodo at https://zenodo.org/records/15421134.

## Notes

https://www2.cloud.editorialmanager.com/pbiology/download.aspx?id=1646396&guid=b1903819-6815-47e2-b93c-fa11d7e11962&scheme=1

### Competing Interest Statement

The authors have declared no competing interest.

### Summary of Updates

This is the Original paper with no modifications made. This Revision includes Supplemental Material at the end, consisting of supplemental figures that are referenced in the paper.

