## Supplemental Figures for "Musical complexity governs a tradeoff between reliability and dimensionality in the neural code"

### Supplementary figures

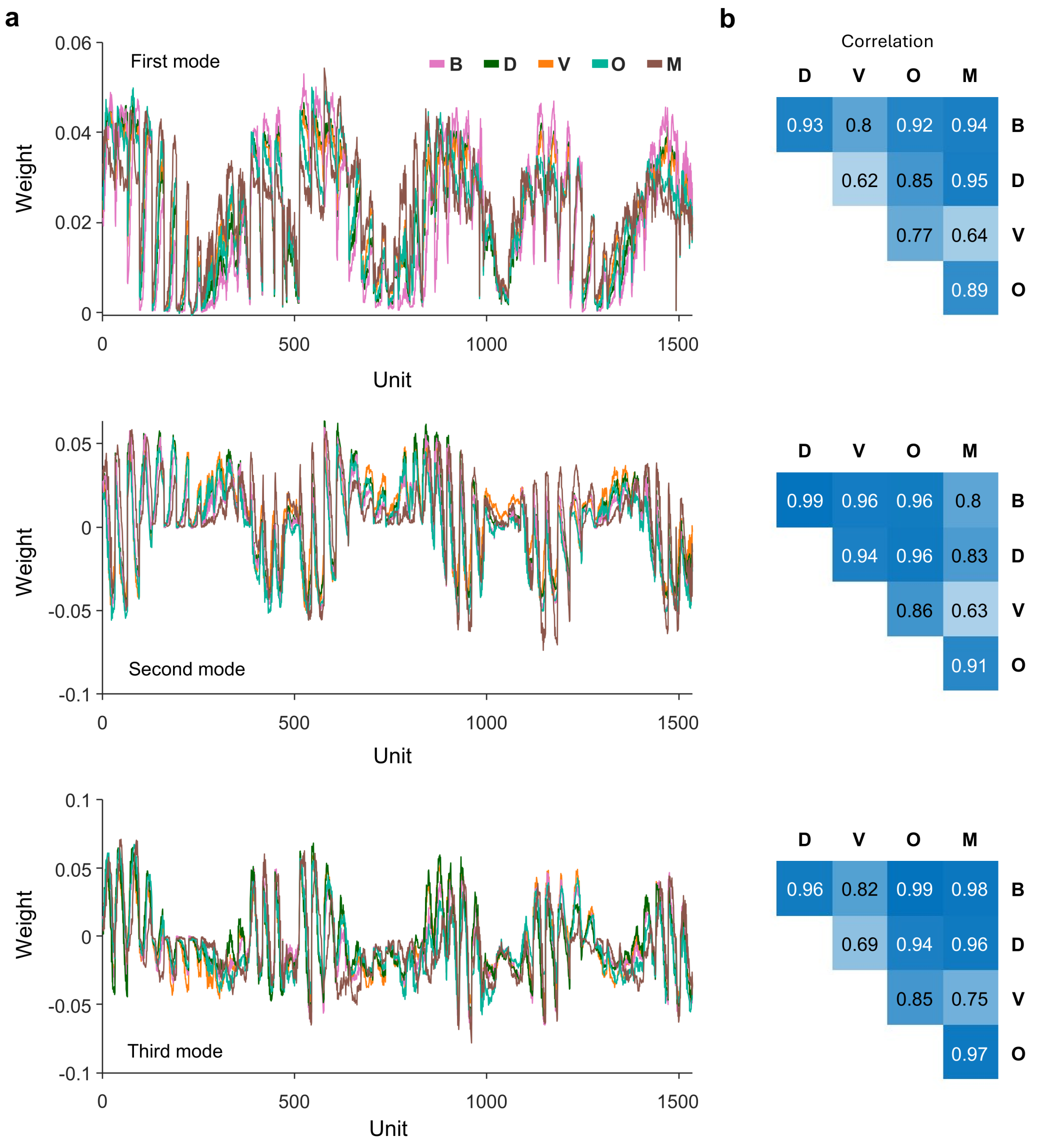


**Figure S1: The leading modes of the signal manifolds for different music types are highly correlated**
**a.** The weights associated with the first three neural modes for each music type for units from animals with normal hearing.
**b.** The correlation between the weights associated with the first three neural modes across music types.

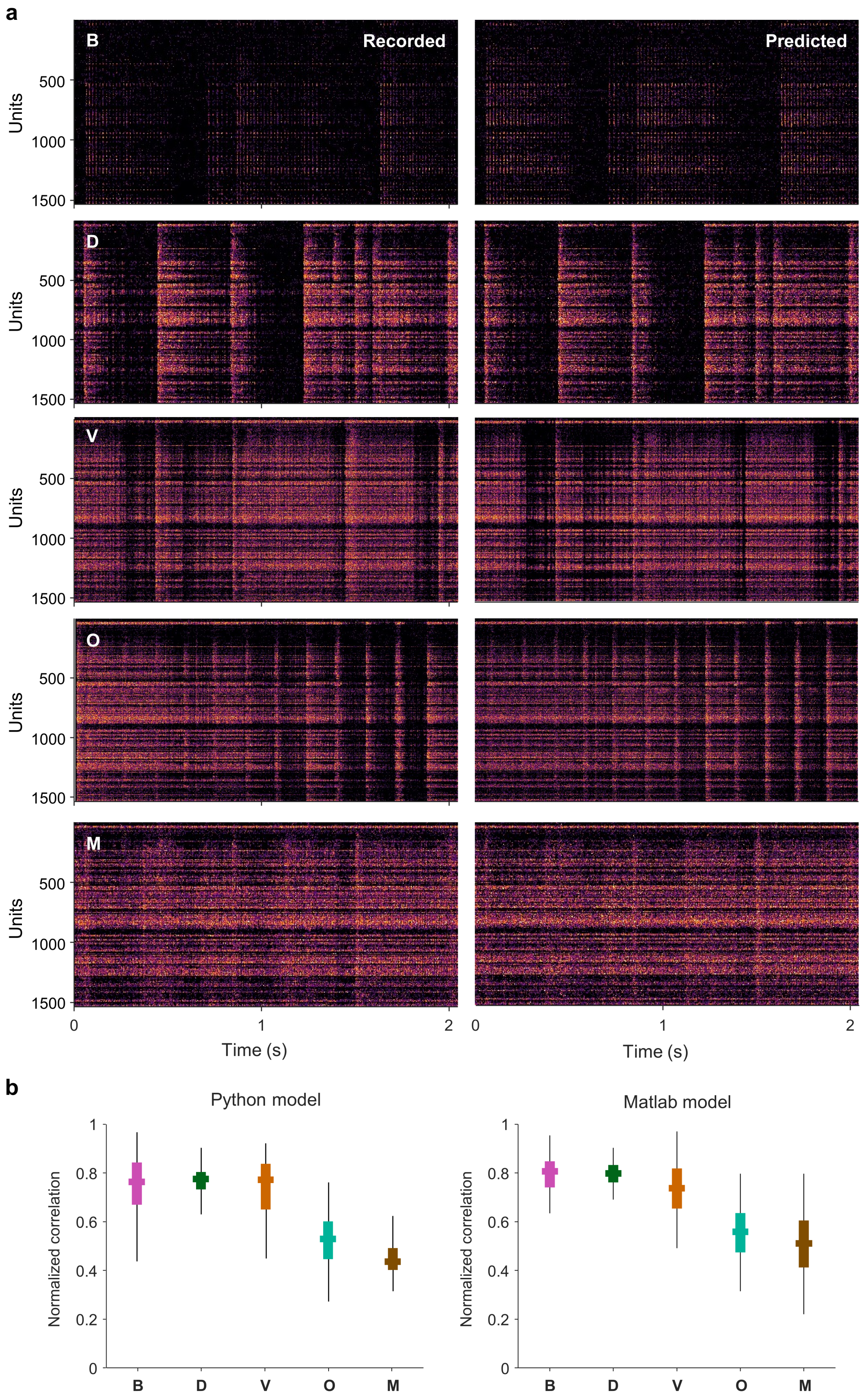


**Figure S2: Examples of recorded and predicted neural activity for different music types**
**a.** Examples of neural responses to music from animals with normal hearing along with the prediction of those responses from the type-specific DNN model with 64 bottleneck channels, with brighter colors indicating more activity and units sorted by preferred frequency (high to low from top to bottom). **b.** The performance of the Python and Matlab versions of the DNN master model (trained on all music). Performance was measured as normalized correlation, the ratio of the correlation between the model prediction and the actual responses to the correlation between the actual responses on repeated trials.

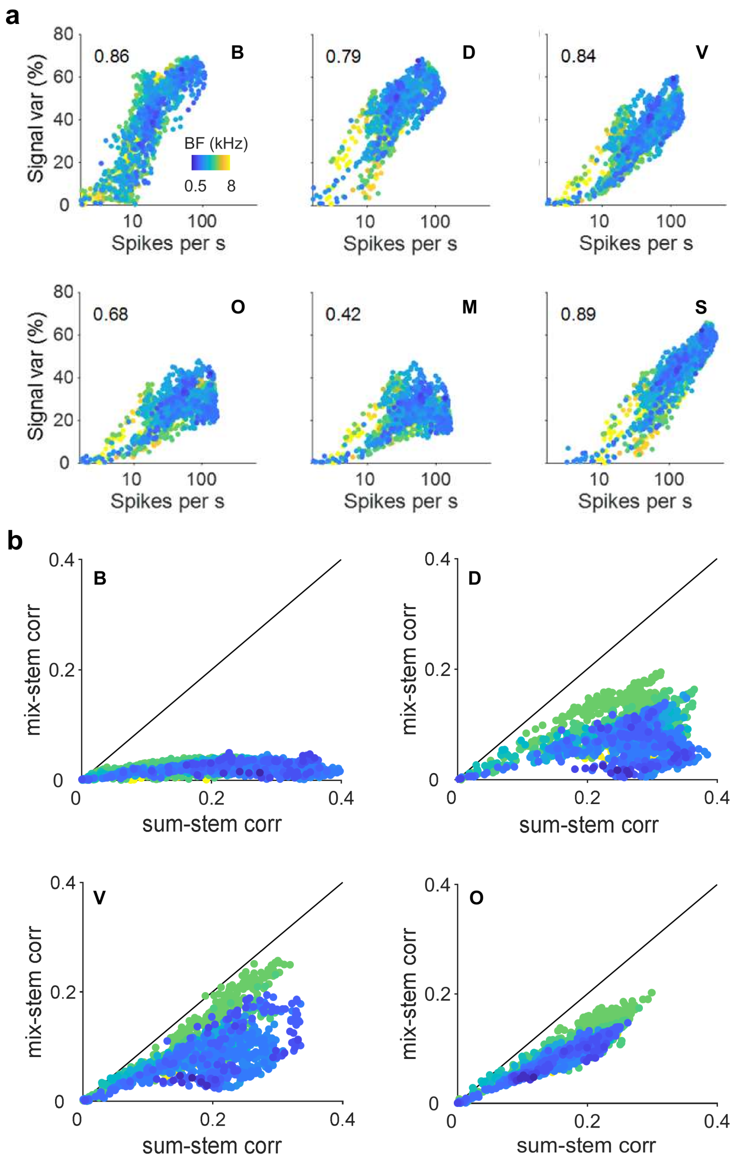


**Figure S3: The properties of neural responses to music for units from animals with hearing loss**
**a.** The strength and reliability of neural responses for all units, with the dots for each unit colored according to its best frequency (analagous to Fig. 1c). **b.** The correlation between stem and mixture responses versus the linear integration benchmark correlation for all units, with the dots for each unit colored according to its best frequency (analogous to Fig. 2d).
